# Patient-derived tumor organoids from resected non-small cell lung cancers for high-throughput response testing with approved and repurposed drugs

**DOI:** 10.1101/2023.10.18.562944

**Authors:** Kanve N. Suvilesh, Yariswamy Manjunath, Yulia I. Nussbaum, Mohamed Gadelkarim, Akhil Srivastava, Guangfu Li, Wesley C. Warren, Chi-Ren Shyu, Feng Gao, Matthew A. Ciorba, Jonathan B. Mitchem, Satyanarayana Rachagani, Jussuf T. Kaifi

## Abstract

**Background:** The five-year survival for non-metastatic non-small cell lung cancer (NSCLC) patients undergoing curative surgery remains poor at ∼50% that is due to locoregional and/or distant metastatic recurrences. Patient-derived tumor organoids (PDTOs) have high potential as clinically relevant high-throughput drug testing platforms to personalize and improve treatment of NSCLC patients. We aimed to develop PDTOs from non-metastatic NSCLC patients to assess their suitability to study tumor heterogeneities and personalized drug responses.

**Methods:** Ten non-metastatic (stage I-IIIA) NSCLC patients undergoing curative surgical resection were prospectively enrolled. PDTOs were established from resected lung tumor tissues and were compared with matched primary tumors by histopathology, immunohistochemistry, whole exome and whole transcriptome sequencing analysis. PDTO responses to standard of care carboplatin/paclitaxel chemotherapy were determined by measuring organoid growth using bright-field 3D imaging. Transcriptomic differential gene expression analysis identified molecular targets for drug repurposing to overcome chemoresistance.

**Results:** NSCLC PDTOs were successfully generated from all 10 (100%) primary tumors with a median time of 12 days (range 4-16 days). All 10 PDTOs could be grown from cryopreserved tumor tissues or reconstituted from frozen PDTOs (living biobank). PDTOs retained histopathological, immunohistochemical protein expression and mutational landscape of the matched primary tumors. Microenvironment cell population analysis revealed epithelial cell signatures of the PDTOs that matched the patients’ lung tumor tissues. Treatment responses of PDTOs to carboplatin/paclitaxel were determined by growth differences versus vehicle control group. 5/10 (50%) PDTOs were chemo-sensitive, whereas 5/10 (50%) were chemo-resistant. Upregulation of aldo-keto reductases (AKR1B10/15) was observed in chemoresistant PDTOs by differential gene expression analysis and confirmed by real-time PCR and immunohistochemistry in PDTOs and tumor tissues. Epalrestat, an anti-diabetic AKR1B10 inhibitory drug, was repurposed to effectively sensitize PDTOs to carboplatin/paclitaxel.

**Conclusions:** PDTOs can be established from resected NSCLC primary tumor tissues with high success rates and conserve cellular, molecular and genomic characteristics of the matched NSCLC tumors. PDTOs can serve as clinically applicable and relevant personalized drug screening platforms to evaluate the therapeutic efficacy of drugs, including repurposed drugs, to overcome chemoresistance.

## BACKGROUND

Lung cancer is by far the number one cause of cancer deaths worldwide. The most common subtype (>80%) is non-small cell lung cancer (NSCLC) (1). Less than a third of NSCLC patients are diagnosed with non-metastatic disease and can undergo surgical resection (2). However, ∼50% of these curatively treated patients develop cancer recurrences later on (2, 3) that are directly linked to drug-resistant, micrometastatic cancer cells that have shed from the primary tumor into the blood, lymphatic system, and distant organs (4, 5). Except the very early-stage IA, NSCLC patients undergoing surgical resection have been routinely administered adjuvant platinum-based doublet cytotoxic chemotherapy to eradicate micrometastatic disease (2, 3), but this therapy leads only to a ∼5% survival benefit (3, 6). Newer targeted (7) and immune therapies (8, 9) also lack sustained effects due to intrinsic or acquired drug resistance (10) and benefit a small fraction of patients only (11). Identifying personalized druggable targets to overcome drug resistances holds great promise to prevent cancer recurrences to improve outcomes for NSCLC patients (3). Exploring gene networks and molecular pathways for identification of agents that are approved for other cancers or diseases can repurpose drugs to overcome drug resistances (12).

A major obstacle in the development of strategies to predict drug treatment responses has been the inability to reliably culture viable patients’ tissues in a high-throughput setting. Patient-derived xenograft (PDX) mouse tumor models have been used widely to study drug resistance patterns in NSCLC (13, 14, 15). However, PDX models are not suitable for high-throughput drug testing because of low engraftment rate, variable and slow tumor growth rates, and high maintenance (16, 17, 18). Patient-derived tumor organoids (PDTOs) are cancer cells grown *ex vivo* in a self-organizing, three-dimensional (3D) matrix in a time-and cost-effective manner (19). PDTOs allow drug screening and these drug responses match the responses observed in solid cancer patients (13, 20, 21, 22). Since PDTOs can be cryopreserved as living biobanks (23), drug sensitivity testing can also be done at later time points (13, 20, 24).

In the present study, we established PDTOs from surgically resected, non-metastatic NSCLC patients’ tumor tissues and tested PDTOs’ clinical applicability for personalized sensitivity screening for standard-of-care chemotherapy and drug repurposing. NSCLC PDTOs maintained molecular (mutational and transcriptomic) characteristics of the matched surgically resected tumor tissues and were suitable for drug testing within a clinically applicable time. Whole transcriptome analyses identified druggable targets to overcome chemoresistance by repurposing an aldo-keto reductase inhibitor (epalrestat) that has been approved for clinical care of diabetics. Our data demonstrate that PDTOs have potential to shape real-time, precision medicine therapy decisions for NSCLC patients.

## METHODS

### Patient enrollment

The study was approved by the University of Missouri Institutional Review Board (IRB#: 2010166) and conducted in accordance with the Declaration of Helsinki. The study was registered at ClinicalTrials.gov (Identifier: NCT02838836; date of registration: July 20, 2016). Study subjects were prospectively recruited at Ellis Fischel Cancer Center at the University of Missouri between June and September 2021. Written informed consent was obtained. Treatment-naïve NSCLC patients were enrolled, whereas patients with concurrent occurrence of another malignancy were excluded. Staging was performed based on the TNM staging manual of the American Joint Committee on Cancer (AJCC), 8^th^ edition (25). All patients recruited were eligible for surgical resection for non-metastatic, loco-regional stages (I-IIIA) as determined by a multidisciplinary thoracic oncology team. All lung cancer resections were performed by specialty-trained thoracic surgeons in alignment with the International Association for the Study of Lung Cancer (IASLC) and Union for International cancer control (UICC). Clinical and outcome data were gathered by reviewing the hospital electronic medical records from our cancer survivorship program or direct communication with the patients or families or their treating physicians.

### Processing of resected lung tumor tissues for PDTO culture

Fragments of the surgically resected primary NSCLC tumors (5-8 mm^3^) from treatment-naïve NSCLC patients were collected in cold advanced Dulbecco’s Modified Eagle Medium (adDMEM) with 10% fetal bovine serum (FBS) and 1% antibiotics (penicillin/streptomycin). All specimens were processed for organoid culture within 20 min of resection (26). Samples were trimmed to remove adjacent, uninvolved normal lung and necrotic lesions, washed thrice with adDMEM+++ (adDMEM supplemented with HEPES, GlutaMAX™ Supplement, Primocin, Amphotericin B) and were cut into pieces (<1 mm^3^ each) with sterile scissors/scalpels. Tumor pieces were incubated with 5 mg/ml collagenase (Gibco), 0.1 mg/ml DNase-I (Sigma-Aldrich), prepared in adDMEM+++ with 10 μM Y-27632 (ROCK inhibitor, R & D systems) at 37 °C for 1 h (shake at 200 rpm) with intermittent agitations. The reaction was stopped using adDMEM+++ supplemented with 10% FBS (200 x *g* for 3 min at 4 °C) and washed with adDMEM+++ (200 x *g* for 3 min at 4 °C). The cell pellet was resuspended in adDMEM+++ and passed through 70-μm cell strainer (BD Falcon). The strained cells were centrifuged at 200 x *g* for 3 min, and the pellet was washed with organoid adDMEM+++ and centrifuged at 200 x *g* for 3 min at 4 °C. The pellet was resuspended in 1/10^th^ volume of culture medium and one volume of Matrigel (Growth factor-reduced; Corning), mixed gently and seeded with ∼20 μl of cell-Matrigel mixture in the center of each well of a 24-well culture plate. The plate was placed upside down at 37 °C for 20 min to facilitate Matrigel dome formation and supplemented with NSCLC organoid culture medium (adDMEM+++ supplemented with human epidermal growth factor (EGF), human fibroblast growth factor (FGF), human noggin, SB202190, B27, N2 supplement, and Y-27632). The medium was changed every 4 days, and the organoids were passaged once >50% of PDTOs reached ≥100 µm in diameter (typically within 1-2 weeks). PDTOs were characterized, stored as frozen stocks, or used for drug testing. Growth time (time from the day of seeding to the day >50% of PDTOs reached ≥100 µm in diameter) was determined, and PDTOs were split 1:4 and passaged to the next generation. Doubling time, defined as time to subsequent passage once fully grown PDTOs occupied the complete area of the Matrigel dome, was determined. PDTOs were maintained for at least 4 passages prior to storing as frozen stocks. PDTOs from each generation were stored in 10% dimethyl sulfoxide (DMSO) in FBS as frozen stocks for future use, serving as a living biobank of PDTOs for future reconstitution.

### Histopathology and immunohistochemical biomarker expression analysis of PDTOs

Matrigel domes containing organoids were scraped off from the culture plate, spun down and culture supernatant was removed. The organoid pellet was fixed in 4% buffered formalin, embedded in paraffin and 4 μm sections were cut and subjected to hematoxylin & eosin (H&E) staining to determine the structural architecture of PDTOs. Additional serial sections were used to characterize the expression of pathological markers by immunohistochemistry (IHC). A small fragment of the matched parental primary tumor tissue was processed in an analogous manner for comparison. Patient-matched primary tumor tissues and PDTOs were immunostained for diagnostic NSCLC markers with antibodies against cytokeratin (CK)7 (mouse, M7018, 1:100; Dako), CK5/6 (mouse, ACR 105, 1:100; Biocare Medical), thyroid transcription factor (TTF-1) (mouse, M3575, 1:100; Dako), and p40 (rabbit, ACI 3030, 1:100; Biocare Medical). Paraffin-embedded tumor tissue sections were processed by de-paraffinization, rehydration, heat-mediated antigen-retrieval, permeabilization and blocking. Tissue sections were incubated with primary antibodies overnight at 4 °C. HRP-conjugated secondary antibodies were incubated for 1 h at room temperature. Sections were treated with DAB for color development and counterstained with hematoxylin (15, 27).

### Drug sensitivity testing in PDTOs

Organoids from passage-2 were trypsinized (TrypLE, Gibco) and mixed in Matrigel and seeded in a 24-well dish in biological triplicates for both vehicle and treatment groups and allowed for 24 h to acclimate. After 24 h (treatment starting time point; Day 0), images from all the wells were taken by brightfield microscopy in a z-stack mode (28, 29). PDTOs were treated with a combination of 1 μg/mL carboplatin and 0.5 μg/mL paclitaxel in organoid culture medium. Doses were determined by titration, as described previously (15). After 72 hours, media was replaced with freshly added chemotherapeutics in organoid medium. All treatments were performed in triplicates, including vehicle controls. Images were taken on days 3 and 6 by z-stack mode in similar focal planes, with day 0 as the baseline for the measurement of growth. On days 0, 3 and 6 (treatment termination), the volume (V) of organoids was calculated (V = 0.5 x Length x Width^2^) (30) from the recorded images. Percentage of organoid growth or regression was determined every 72 h against vehicle control groups, using day 0 as baseline. An average of 12-15 organoids per well were monitored to determine the efficacy of chemotherapy in inhibiting organoid growth. Patient tumors were categorized as resistant or sensitive based on PDTOs therapy outcome. To determine the potential of aldo-keto reductase family 1 member B10 (AKR1B10) inhibitor to overcome the chemotherapy resistance to carboplatin and paclitaxel, resistant PDTOs were treated with chemotherapeutics in presence or absence of 1 µM epalrestat (SML0527; Sigma-Aldrich). Dose of epalrestat was determined following generation of a cytotoxicity dose response curve using PDTOs. To further confirm the chemoresistance reversal by epalrestat, PDTO Matrigel domes from different treatment groups were fixed in 4% buffered formalin and processed for IHC staining, as mentioned above, for the proliferation marker Ki67 (rabbit, ab16667, 1:200; Abcam) and apoptosis marker cleaved caspase-3 (Rabbit, 9664, 1:100; Cell Signaling Technologies). Percentages of Ki67 and or cleaved caspase-3 positive expression in PDTOs were calculated by a pathologist blinded to the study. Matched lung tumor tissues from chemoresistant and chemosensitive patients (as determined in PDTOs) were immunostained with anti-AKR1B10 antibody (rabbit, ab192865; 1:1000; Abcam) and detected by DAB method.

### Whole exome sequencing of PDTOs and matched parental primary lung tumor tissues

*DNA extraction:* PDTOs from passages 1-3 were cultured to confluence and incubated with Cell Recovery Solution (Corning) for 30 min at 4° C, followed by washing (x3) with ice cold PBS to remove Matrigel. PDTO pellets and matched primary lung tumors (∼10mg tissue) were used for genomic DNA extraction using DNeasy Blood and Tissue kit (Qiagen). Genomic DNA was eluted with buffer AE (supplied in the kit) and diluted to a concentration of 100 ng/µL for library preparation. Integrity of extracted genomic DNA was confirmed by agarose gel electrophoresis.

*Library Preparation, Sequencing:* The genomic DNA was randomly sheared into short fragments with the size of 180-280 bp. The obtained fragments were end repaired, A-tailed, and further ligated with Illumina adapters. The fragments with adapters were PCR amplified, size selected, and purified. Hybridization capture of libraries was proceeded according to the following procedures. Briefly, the libraries were hybridized in the buffer with biotin-labeled probes, and magnetic beads with streptavidin were used to capture the exons of genes. Subsequently, non-hybridized fragments were washed out and probes were digested. The captured libraries were enriched by PCR amplification. The library was quantified with Qubit, real-time PCR and a bioanalyzer for size distribution detection. Quantified libraries were pooled and sequenced on Illumina platforms with PE150 strategy, to 30X coverage depth.

### Bioinformatics Analysis

#### Quality Control and alignment

To remove low-quality reads and sequences pair of reads with adapter contamination, pair of reads having more than 10% uncertain bases and with low quality Phred score were discarded. Clean reads were mapped to the reference genome (b37/hg19/hg38) by Burrows-Wheeler Aligner (BWA) software (31) to generate BAM files. Subsequently, Sambamba (32) was used to sort BAM files according to chromosome position. Picard tools were then utilized to merge BAM files and mark duplicate reads.

#### Variant Calling and Annotation

GATK (33) was used to call SNP and InDel and ANNOVAR (34) was used to perform variant annotation. Further to find cancer-relevant somatic mutations and categorize identified mutations to tier 1 and tier 2, variants were filtered against somatic mutations reported in Catalogue Of Somatic Mutations In Cancer (COSMIC) database (35) and COSMIC Cancer Gene Census tier 1 and tier 2 curated mutations (36), respectively.

#### Whole transcriptome analysis of PDTOs and matched primary lung tumor tissues

*RNA extraction*: To determine global transcriptomic signatures, total RNA from PDTOs and matched primary lung tumors was extracted using RNeasy kit (Qiagen) and solubilized in nuclease free water at a concentration of 100 ng/µL for library preparation. *Illumina Stranded Total RNA Library Preparation and Sequencing:* Libraries were constructed following the manufacturer’s protocol with reagents supplied in Illumina’s Ribo Zero Gold rRNA Removal Kit (Human/Mouse/Rat) and Illumina Stranded RNA Prep, Ligation Kit. Prior to rRNA depletion, the sample concentration was determined by Qubit fluorometer (Invitrogen) using the Qubit HS RNA assay kit, and the RNA integrity was confirmed using the Fragment Analyzer automated electrophoresis system. Briefly, the rRNA is removed from total RNA by hybridization probes using the Ribo Zero kit. The recovered mRNA is fragmented, and double-stranded cDNA is generated. Pre-index anchors are ligated to the ends of cDNA. A subsequent PCR step is done to selectively amplify the anchor-ligated DNA fragments and add indexes and primer sequences for cluster generation. The final amplified cDNA constructs were purified by addition of Axyprep Mag PCR Clean-up beads. The final construct of each purified library was evaluated using the Fragment Analyzer automated electrophoresis system, quantified with the Qubit fluorometer using the Qubit HS dsDNA assay kit, and diluted according to Illumina’s standard sequencing protocol for sequencing on the NovaSeq 6000. High-throughput sequencing was performed at the University of Missouri Genomics Technology Core.

#### RNA-seq analysis

Quality control of RNA-seq fastq files was performed using MultiQC tool (37). Fastq files of RNA-seq data from 20 samples were aligned to human reference genome (GRCh38) using STAR alignment method (38). Based on alignment and quality control results, one tumor sample and the matched PDTO (MU369) were removed from further analyses. Reads were quantified by featureCounts algorithm into count matrices (39). To find differentially expressed genes between samples that were treatment resistant and responsive, we used DESeq2 R package with specification of Wald test (40). To quantify cellular populations in each sample, we used Microenvironment Cell Populations-counter (MCP-counter) method (41). For estimating epithelial cells’ subpopulations customized list of cell specific markers was added to MCP-counter gene list (**Supplementary File 1**). Epithelial cell specific markers were selected from the Lung Gene Expression Analysis (LGEA) portal (42), The Human Protein Atlas (43) and Cell marker database (44). Principal component analysis (PCA) plot was built to determine degree of similarity or variability between different types of models such as patient primary tumor samples, PDTOs, and patient-derived xenografts (PDXs) from an internally generated “Biobank” (15). Briefly, data from 9 patient primary tumor samples, 9 PDTOs, and 4 PDX tumors were projected onto the first two principal components using plotPCA function of DESeq2 R package. Before plotting, counts were normalized and applied variance stabilizing transformation to UMI count data using a regularized Negative Binomial regression model.

### Real-time PCR (qPCR)

To validate the expression of AKR1B10 observed in differential gene expression analysis, qPCR was performed. Reverse transcription of total RNA to cDNA was conducted with a High-Capacity cDNA Reverse Transcription Kit (Applied Biosystems). Primer sets were designed using NCBI primer pick website, which considers sequence alignments, primer stability, hairpins, primer melting temperatures, and primer-dimer interactions. Primer sets were selected to have a melting temperature between 58-60^0^C and guanine-cytosine content ∼50% and to generate a PCR amplicon <100-300 bp. BLAST (Basic Local Alignment Search Tool) was used to check for uniqueness, including the presence of amplifiable pseudogenes (45). All primers were synthesized by Sigma-Aldrich [sequences: AKR1B10-Forward (5’-3’) CAGAAGACCCTTCCCTGCTG, Reverse (5’-3’) CGTTACAGGCCCTCCAGTTT, GAPDH-Forward (5’-3’) TTCTTTTGCGTCGCCAGCC, Reverse (5’-3’) TCCCGTTCTCAGCCTTGACG]. qPCR was performed with QuantStudio 3 Detection System (Thermo Fisher Scientific) in a 20 μL reaction mixture containing SYBR Green PCR Master Mix (Thermo Fisher Scientific). Reactions were run in triplicates. The expression of AKR1B10 was normalized to the geometric mean of housekeeping gene GAPDH to control the variability of expression levels. The data were analyzed using the 2^−ΔΔCT^ method and the relative fold change of AKR1B10 in chemoresistant patient tumors against chemosensitive patient tumors was determined.

### Statistical analyses

Statistical analyses involving multiple experimental groups were performed using one-way ANOVA (Kruskal-Wallis Multiple Comparison and Tukey’s multiple comparisons test) and Student’s t-test (Multiple t-tests). Differentially expressed genes were identified by Wald test. Significance statements refer to a *p* value of <0.05. Statistical analyses were performed using software Prism version 8.00 (GraphPad, La Jolla, CA).

## RESULTS

### Clinicopathological characteristics of NSCLC patients enrolled for PDTO generation

Ten treatment-naïve NSCLC patients undergoing curative surgical resection for non-metastatic cancers were prospectively recruited. Clinical details of the enrolled NSCLC patients are described in **Table 1**. Five (50%) patients were females. The median age for male patients was 63 years (range 59-71) and for female patients 64 years (57–67). Seven (70%) patients had stage I per AJCC criteria. One (10%) patient had stage II, and two (20%) patients were incidentally diagnosed with stage IIIA by identification of cancer-positive mediastinal (N2) lymph nodes in the pathologic specimen from the surgical resection. Further, five patients (50%) had lung squamous carcinoma (LUSC) and lung adenocarcinoma (LUAC), respectively. Furthermore, three (30%) patients had histological grade III tumors, whereas 7 patients (70%) had grade II tumors. Following surgical resection, the post-operative surveillance period for patients who remained recurrence-free and alive was a median of 25 months (range 24-26).

During this observation period, only one (10%) patient (MU383) developed a cancer recurrence and passed away 15 months after surgery due to progression of his metastatic disease that he developed while receiving adjuvant chemotherapy (carboplatin/paclitaxel) two months after his surgery, indicating chemoresistance. Notably and as outlined below, the PDTO derived from this patient also showed chemoresistance towards carboplatin/paclitaxel.

**Table 1.**
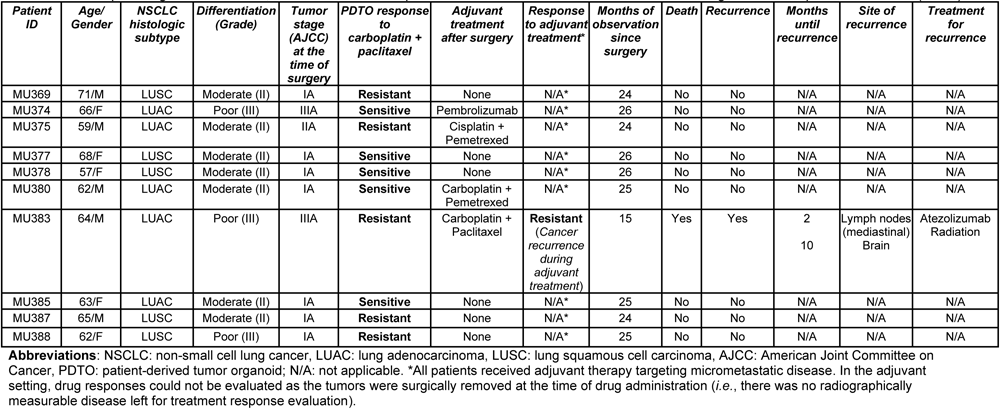
Clinicopathological information, PDTO treatment responses and recurrence/survival status of the surgical NSCLC patients enrolled (N=10).

| *Patient ID* | *Age/*<br>*Gender* | *NSCLC*<br>*histologic*<br>*subtype* | *Differentiation*<br>*(Grade)* | *Tumor*<br>*stage*<br>*(AJCC)*<br>*at the*<br>*time of*<br>*surgery* | *PDTO response*<br>*to*<br>*carboplatin +*<br>*paclitaxel* | *Adjuvant*<br>*treatment*<br>*after surgery* | *Response*<br>*to adjuvant*<br>*treatment\** | *Months of*<br>*observation*<br>*since*<br>*surgery* | *Death* | *Recurrence* | *Months*<br>*until*<br>*recurrence* | *Site of*<br>*recurrence* | *Treatment*<br>*for*<br>*recurrence* |
| --- | --- | --- | --- | --- | --- | --- | --- | --- | --- | --- | --- | --- | --- |
| MU369 | 71/M | LUSC | Moderate (II) | IA | **Resistant** | None | N/A* | 24 | No | No | N/A | N/A | N/A |
| MU374 | 66/F | LUAC | Poor (III) | IIIA | **Sensitive** | Pembrolizumab | N/A* | 26 | No | No | N/A | N/A | N/A |
| MU375 | 59/M | LUAC | Moderate (II) | IIA | **Resistant** | Cisplatin +<br>Pemetrexed | N/A* | 24 | No | No | N/A | N/A | N/A |
| MU377 | 68/F | LUSC | Moderate (II) | IA | **Sensitive** | None | N/A* | 26 | No | No | N/A | N/A | N/A |
| MU378 | 57/F | LUSC | Moderate (II) | IA | **Sensitive** | None | N/A* | 26 | No | No | N/A | N/A | N/A |
| MU380 | 62/M | LUAC | Moderate (II) | IA | **Sensitive** | Carboplatin +<br>Pemetrexed | N/A* | 25 | No | No | N/A | N/A | N/A |
| MU383 | 64/M | LUAC | Poor (III) | IIIA | **Resistant** | Carboplatin +<br>Paclitaxel | **Resistant**<br>(Cancer<br>recurrence<br>during<br>adjuvant<br>treatment) | 15 | Yes | Yes | 2<br>10 | Lymph nodes<br>(mediastinal)<br>Brain | Atezolizumab<br>Radiation |
| MU385 | 63/F | LUAC | Moderate (II) | IA | **Sensitive** | None | N/A* | 25 | No | No | N/A | N/A | N/A |
| MU387 | 65/M | LUSC | Moderate (II) | IA | **Resistant** | None | N/A* | 24 | No | No | N/A | N/A | N/A |
| MU388 | 62/F | LUSC | Poor (III) | IA | **Resistant** | None | N/A* | 25 | No | No | N/A | N/A | N/A |

**Abbreviations:** NSCLC: non-small cell lung cancer, LUAC: lung adenocarcinoma, LUSC: lung squamous cell carcinoma, AJCC: American Joint Committee on Cancer, PDTO: patient-derived tumor organoid; N/A: not applicable. \*All patients received adjuvant therapy targeting micrometastatic disease. In the adjuvant setting, drug responses could not be evaluated as the tumors were surgically removed at the time of drug administration (*i.e.*, there was no radiographically measurable disease left for treatment response evaluation).

### PDTOs have heterogeneous growth patterns and can be reconstituted from frozen stocks

All 10 (100%) freshly resected primary NSCLC tumor tissues were successfully grown to PDTOs (**Fig. 1A**). PDTOs were passaged into subsequent generations after >50% of PDTOs grew to a size of ∼100 μm (**Fig. 1B**). PDTO growths were observed daily using 3D (z-stack) imaging using bright-field microscopy (**Fig. 1C**). Comparison showed heterogeneous PDTO growth rates with a median of 12 days (range 4-16 days) and with a median doubling time of 7 days (range 5-13 days) (**Fig. 1B**). All 10 (100%) PDTOs could also be successfully established from cryopreserved primary lung tumor tissues or reconstituted from cryopreserved frozen PDTOs, providing a “living biobank” resource for later study.

**Figure 1.**
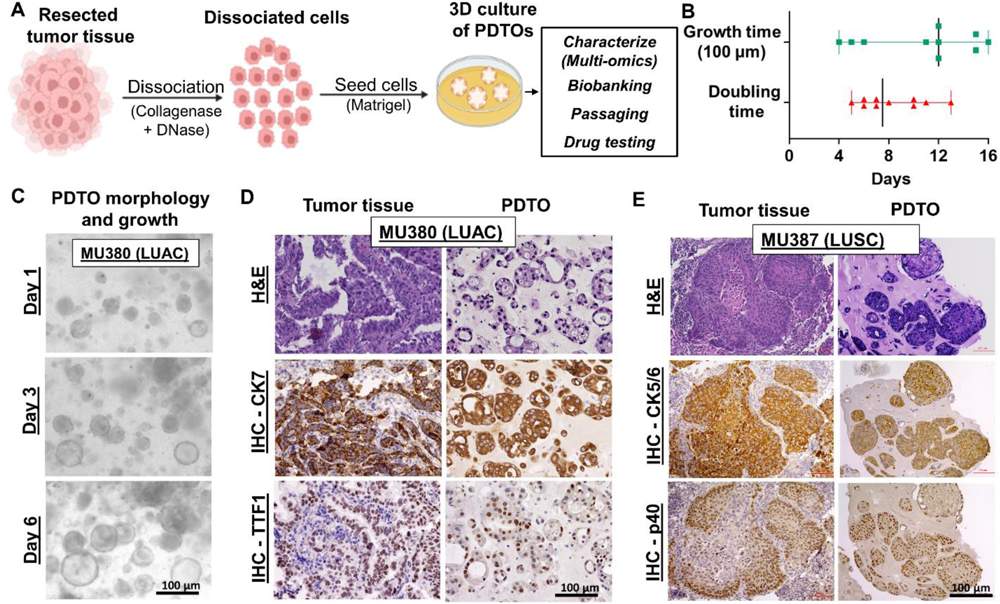
Non-small cell lung cancer (NSCLC) patient-derived tumor organoid (PDTO) development and characterization. **A:** An outline depicting development and characterization of PDTOs. Resected primary NSCLC tumors were dissociated, seeded with Matrigel, and cultured in organoid medium to generate PDTOs that were used for different analyses. **B:** Time for organoids to grow to 100 µm diameter from the day of organoid seeding (growth time) and doubling time (defined as time the first passage of PDTOs took to occupy the complete area of the Matrigel dome requiring transfer to second passage). **C:** Morphology by bright-field microscopy of a representative PDTO (patient MU380) on days 1, 3 and 6. (Magnification: 20x; scale bar, 100 µm). **D and E:** Comparison of PDTOs and matched primary tumor tissues demonstrating conservation in histomorphology (hematoxylin and eosin; upper panels), and pathological biomarker expression (immunohistochemistry (IHC); middle and lower panels). Representative **D:** Patient MU380: Lung adenocarcinoma (LUAC) - specific markers [cytokeratin (CK) 7 and thyroid transcription factor (TTF1)], and representative **E:** Patient MU 387: Lung squamous cell carcinoma (LUSC) - specific markers (CK5/6 and p40) were consistently expressed in PDTOs and matched primary tumor tissues. (Magnification: 20x; scale bar, 100 µm).

### PDTOs retain histopathological characteristics and NSCLC biomarker expression

Formalin-fixed, paraffin-embedded PDTOs were stained with H&E to compare the histopathology of PDTOs with matched primary tumors. We observed that the histological features of primary tumors were preserved in PDTOs. The LUAC-derived PDTOs conserved tumor cell architecture with typical glandular appearance (*e.g.*, cystic or acinar structures) like the matched primary tumors (**Fig. 1D; upper panels**). LUSC-derived PDTOs retained characteristic solid polygonal cells with nested growth patterns and intercellular bridges (lacking glandular structure) with hyperchromatic nuclei, as observed histologically in matched primary tumors (**Fig. 1E; upper panels**). Further characterization was performed by IHC staining of LUAC-and LUSC-specific diagnostic pathological markers in patient-matched PDTOs and tumor tissues. PDTOs conserved diagnostic pathological biomarker [CK7 and TTF-1 in LUAC-PDTOs (**Fig. 1D; middle and lower panels**), CK5/6 and P40 in LUSC-PDTOs (**Fig. 1E; middle and lower panels**)] expressions that were also detected in matched primary tumors. Taken together, all ten PDTOs consistently retained the tumor architecture and biomarker expression of matched primary NSCLC tumor counterparts.

### PDTOs recapitulate the genomic fidelity of the matched parental NSCLC tumors

Mutational signatures of six matched pairs of NSCLC PDTOs and primary tumors were determined by whole exome sequencing (**Fig. 2**). Hierarchical clustering based on somatic mutations demonstrated strong similarities between matched NSCLC primary tumors and PDTOs (**Fig. 2A**). Additionally, striking overlap of cancer-relevant somatic mutations (reported in COSMIC) such as missense, nonsense, frameshift deletion and insertion mutations confirmed conserved genetic polymorphisms between NSCLC primary tumors and matched PDTOs (**Fig. 2B** **and Supplementary File 1 and 2**). Moreover, somatic mutations identified in oncogenes and tumor suppressor genes were also overlapped between primary tumors and PDTOs (**Fig. 2C** **and Supplementary File 1 and 2**). Based on the COSMIC database cancer gene census, we also categorized cancer-relevant mutations hierarchically in pan-cancer and NSCLC tier 1 and tier 2, respectively (total of four categories), again observing broad overlap in these groups (**Fig. 2D**). Finally, heatmap of specific and most relevant pan-cancer and NSCLC somatic mutations identified in oncogenes and tumor suppressor genes is shown in **Fig. 2E**, demonstrating vast overlap between the matched pairs except in one case where a KRAS G12V mutation was not observed in the matched PDTO (MU374) (Table S1). In summary, whole exome sequencing analysis demonstrated that NSCLC PDTOs maintained the genomic fidelity as determined by genetic polymorphisms and genomic heterogeneity of their original lung tumor tissues.

**Figure 2.**
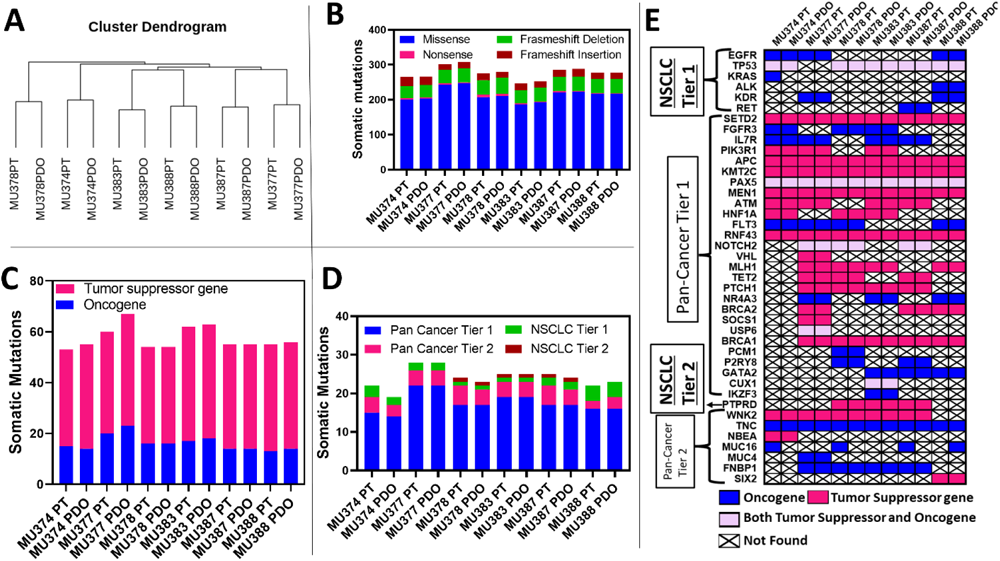
Mutational signatures in matched NSCLC patient-derived tumor organoids (PDTOs) and parental primary lung tumor tissues (PTs). Whole exome sequencing was performed in matched PDTOs and PTs (N=6 matched pairs). **A:** Hierarchical clustering and dendrogram predicting similarities and variabilities between PDTOs and PTs based on somatic mutations**. B:** Total number of cancer-relevant somatic mutations identified in PTs and matched PDTOs (somatic mutations were identified using COSMIC database). **C:** Total number of somatic mutations identified in tumor suppressor genes and oncogenes. **D:** Total number of Pan-Cancer Tier 1 and 2, NSCLC Tier 1 and Tier 2 mutations identified in PTs and matched PDTOs. Tier 1 and Tier 2 mutations were screened using COSMIC cancer gene census. **E:** Heatmap showing somatic mutations in oncogenes and tumor suppressor genes identified in PTs and matched PDTOs.

### Epithelial cancer cell heterogeneity is retained in NSCLC PDTOs

Whole transcriptome analysis by using total RNA from primary tumors and matched PDTOs was performed to determine the global gene expression profiles and cellular heterogeneity. For additional comparison, an internal library of NSCLC patient tumors-derived xenograft (PDX) tumors grown in immunodeficient mice were also included in the principal component analysis (PCA). PCA revealed that tumor tissues and PDTOs were closely clustered in comparison to PDX tumors with more similarities in gene expression between tumor tissues and PDTOs in contrast to PDX tumors (**Fig. 3A**). Cell type abundance was determined using microenvironment cell population (MCP-counter) method which revealed conserved epithelial cell composition in all PDTOs compared to matched primary NSCLC tumor tissues for both LUAC and LUSC, except MU380 (**Fig. 3B**; **Fig. S1**). However, heterogeneity with respect to conservation of immune, stromal, and endothelial cell types was not universally observed. For instance, immune, stromal, and endothelial along with epithelial cell types were conserved only in MU378 and MU385 patient matched tumor tissues and PDTOs, accounting for only ∼20% of all the samples (**Fig. 3B, right panel**; **Fig. S1**). In contrast, these cell types were not conserved in MU377-derived PDTOs as well as other PDTOs (**Fig. 3B, left panel;** **Fig. S1**). Epithelial cells were identified with cell-specific markers (**Supplementary File 1**). As expected, epithelial subtype signatures confirmed the presence of alveolar epithelial cell type II (AT-2) in LUAC-derived PDTOs and matched primary tumor tissues (**Fig. 3C, left panel**). Similarly, basal cells were abundantly present in LUSC-derived PDTOs and matched tumor tissues (**Fig. 3C, right panel**). In contrast to other cell types, LUAC and LUSC subtype-specific epithelial cell signatures were consistently conserved in PDTOs.

**Figure 3.**
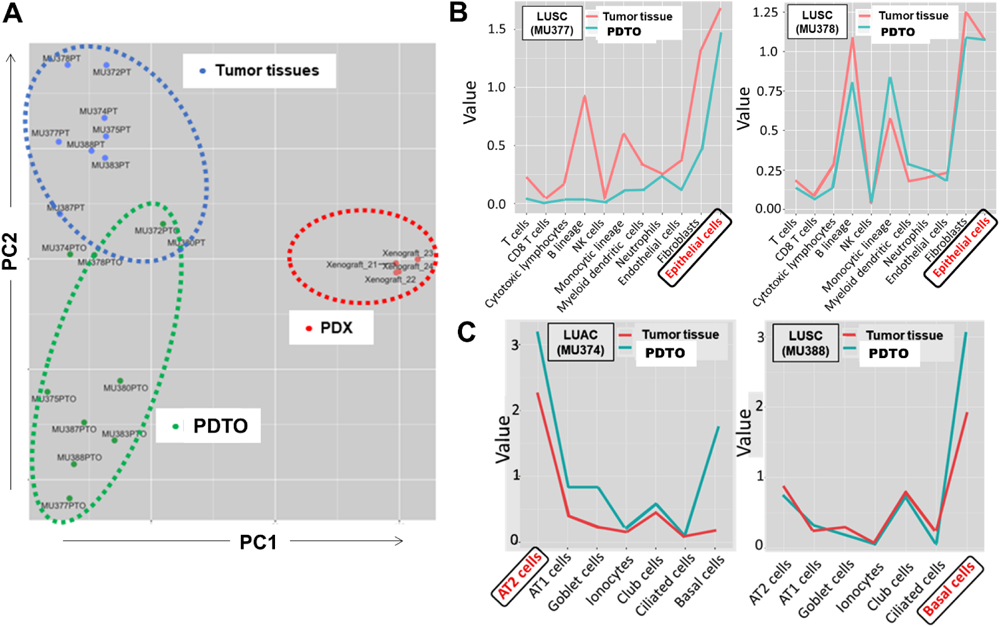
Gene expression and cellular heterogeneity comparison between PDTOs and matched patient primary lung tumor tissues (PT). **A:** Principal component analysis (PCA) of PTs (N=9), PDTOs (N=9) and an internal library of patient-derived NSCLC xenograft (PDX) mouse models (N=4) whole transcriptome sequencing (bulk RNA-seq) analyses, demonstrating that PDTOs and PTs clustered closer to each other than PDX tumors grown in immunodeficient mice. **B:** Microenvironment Cell Populations (MCP)-counter graphs showing cellular abundance of PDTOs and matched PTs and of two representative NSCLC patients with conserved epithelial cell populations, whereas immune cell compositions were found to be inconsistently preserved [preserved in patient MU378 (right panel), not entirely preserved in patient MU377 (left panel)]. **C:** Epithelial cell type abundance is conserved regarding AT2 cells in a LUAC (MU374; left panel), and basal cell abundance in a LUSC (MU388; right panel) in PDTOs and matched PTs, respectively.

### PDTOs serve as high-throughput drug response testing platforms

To determine the clinical applicability of PDTOs as drug screening platforms, organoids were treated with a standard-of-care platinum-based doublet chemotherapy (carboplatin/paclitaxel) regimen (**Fig. 4A**). Responses to chemotherapy in PDTOs were quantified based on growth changes on days 3 and 6 using real-time 3D optical z-stack imaging in comparison to vehicle control. PDTOs were categorized as chemosensitive in cases of statistically significant (p<0.05; Student’s t-test) reduction of PDTO growth, and chemoresistant if growth was maintained in presence of carboplatin/paclitaxel (**Fig. 4B-K**; **Table 1**). We observed heterogeneous chemoresponses with 5 (50%) PDTOs sensitive and 5 (50%) PDTOs resistant to carboplatin/paclitaxel (**Fig. 4**; **Table 1**), consistent with the ∼50% chemoresistance rates typically observed in NSCLC patients. Of note, the observed chemoresponses of the PDTOs assessed at the two measurement time points on day 3 and 6 were equivalent in all PDTOs. Patient-matched chemoresistance was observed in patient MU383 who had tumor recurrences while being under treatment with carboplatin/paclitaxel, consistent with chemoresistance observed in patient MU383-derived tumor organoid (**Table 1**).

**Figure 4.**
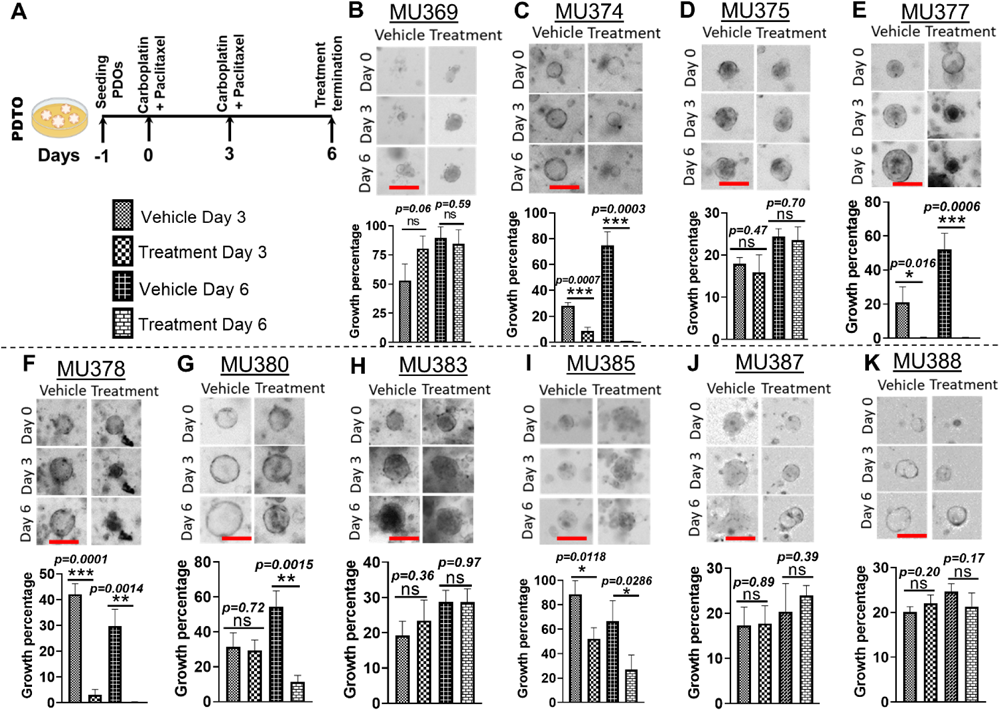
Patient-derived tumor organoids as drug testing platforms for standard-of-care chemotherapy. **A:** Diagram showing treatment protocol. **B-K:** Growth comparisons of PDTOs in chemotherapy treatment (carboplatin + paclitaxel) vs. vehicle control groups, **Upper panels:** Representative bright-field microscope images of PDTOs from vehicle and carboplatin + paclitaxel treatment groups in ten NSCLC patients tested (scale bar – 200 μm). **Lower panels:** Growth percentages of PDTOs in control and treatment groups on day 3 and day 6 in comparison to baseline on day 0. Data are presented as mean ± standard error of the mean (SEM). Statistical analysis was performed using Students *t*-test. ***-*p*<0.001; **- *p*<0.005; *- *p*<0.05; ns- Not significant.

### Differential gene expression analysis identified targets for drug repurposing in chemoresistant PDTOs

Differential gene expression analysis showed significant upregulation of genes (AKR1B10, AKR1B15, HORMAD1, RPL17P11, S100A7/8, RF00003) in chemoresistant PDTOs, indicating a potential role of these genes in chemoresistance (**Fig. 5A**; **Supplementary File 4**). Among differentially upregulated genes in chemoresistant PDTOs, aldo-keto reductase family genes (AKR1B10 and AKR1B15) have been reported to have a role in detoxification of cytotoxic compounds leading to induction of platinum or taxol-based drug resistance. As targetable gene to overcome resistance to carboplatin/paclitaxel, AKR1B10 was chosen for further study. For validation, matched patient primary lung tumor tissues were categorized into resistant or sensitive based on PDTO chemotherapy outcome and AKR1B10 expression was determined by differential gene expression analysis. As observed in chemoresistant PDTOs, matched patient tumors showed overexpression of AKR1B10 (**Fig.5G**; **Supplementary File 5**).

**Figure 5.**
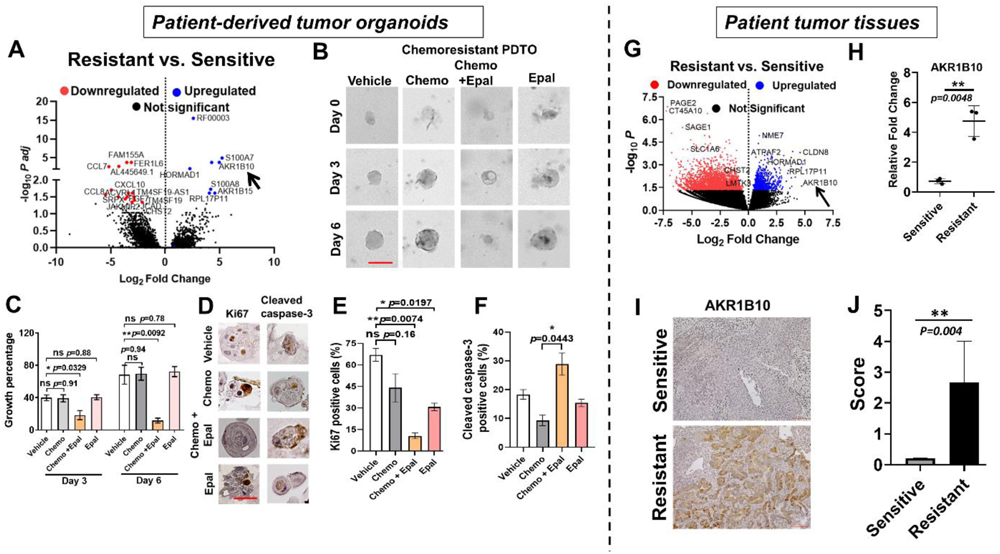
Differentially expressed genes in chemoresistant vs. chemosensitive patient-derived tumor organoids (PDTOs) and reversal of chemoresistance by drug repurposing. **A:** Differential gene expression analysis of chemosensitive (N=4) vs. chemoresistant (N=4) NSCLC PDTOs treated with carboplatin/paclitaxel (arrow: AKR1B10). **B:** Overcoming chemoresistance by repurposing an FDA-approved aldoketoreductase inhibitor (epalrestat). Representative images of chemoresistant PDTOs (day 0, 3 and 6) treated with chemo (carboplatin/paclitaxel), chemotherapy plus epalrestat (Scale bar - 200μm). **C:** Growth percentages of PDTOs in different treatment groups determined on day 3 and day 6 (relative to day 0). Growth measurements were performed in triplicates and the data are represented as mean ± SEM. Statistical analysis performed using one-way ANOVA and *p* value was determined by Tukey’s multiple comparisons test; * - *p*<0.05. **-*P*<0.01. **D:** Proliferation is decreased, and apoptosis is increased in chemoresistant PDTOs treated with epalrestat. PDTOs were harvested on day 6 and analyzed by immunostaining for proliferation marker (Ki67) and apoptosis marker (Cleaved caspase 3). Representative brightfield microscope images are presented. (Scale bar - 200μm). **E and F:** Percentages of Ki67-positive **(E)** and cleaved caspase 3-positive **(F)** cells from different treatment groups are represented as mean ± SEM (Kruskal-Wallis Multiple Comparison; * - P<0.05, **-*P*<0.01). Right Panel- Validation of AKR1B10 expression in chemoresistant versus chemosensitive patient tumors (categorized based on PDTOs therapy outcome). **G:** Differential gene expression analysis revealed overexpression of AKR1B10 in patient primary lung tumor tissues (PTs) similar to expressions in PDTOs that were found to be chemoresistant. **H:** qPCR measurement of mRNA expression of AKR1B10 gene in chemosensitive (n=4) and chemoresistant (n=4) PTs. **I:** Immunostaining of AKR1B10 in chemosensitive (n=4) and chemoresistant (n=4) PTs showing high expression in chemoresistant tumors. **J:** Blinded scoring of AKR1B10 expression assessed by immunostaining between chemoresistant and -sensitive data are presented as mean ± SD. Statistical analysis was performed using Student’s t test; * - *p*<0.05. **-*p*<0.005.

Moreover, AKR1B10 expression by qPCR was found to be >5-fold higher in resistant patient tumors relative to sensitive tumors (p=0.0048; Student’s *t*-test) (**Fig. 5H**). In addition, chemoresistant tumors have significantly higher AKR1B10 protein expression compared to chemosensitive tumors as determined by IHC (**Fig. 5I**) and blinded scoring (**Fig. 5J**). For reversal of chemoresistance, an inhibitor of AKR1B10 - epalrestat - that is already in clinical use for patients with diabetic polyneuropathy was selected in this study. Doses of epalrestat for drug repurposing experiments were selected based on dose titration curves against PDTOs (**Fig. S2 A** **and** **B**).

### Overcoming chemoresistance with the repurposed AKR1B10 inhibitor drug epalrestat

To evaluate the therapeutic potential of epalrestat in reversal of chemoresistance against carboplatin/paclitaxel, chemoresistant PDTOs were reconstituted from the frozen living biobank. Treatment with epalrestat significantly sensitized PDTOs to carboplatin/paclitaxel as evidenced by the inhibition of PDTO growth compared to the persistent growth of PDTOs treated with carboplatin/paclitaxel alone (**Fig. 5** **B and C**) (p<0.05; one-way ANOVA and Tukey’s multiple comparisons test; **Fig. 5** **B and C**). Epalrestat alone did not have any significant effects on PDTO growth (**Fig. 5** **B and C**). To further confirm cytotoxicity induced by carboplatin and paclitaxel in presence of epalrestat, we checked the expression of cell proliferation and apoptosis markers. IHC for cell proliferation (Ki67) and apoptosis (cleaved caspase-3) marker expression revealed a significant reduction in number of proliferating (Ki67-positive) cells (**Fig. 5D, left panels;** **Fig. 5E**) and increased number of apoptotic (cleaved caspase-3-positive) cells (**Fig. 5D, right panels;** **Fig. 5F**) in PDTOs treated with epalrestat and carboplatin/paclitaxel combination compared to other groups (p<0.05; Kruskal-Wallis multiple comparison test). These results indicate that NSCLC PDTO chemoresistance is associated with upregulated aldo-keto reductase expression that can be overcome by repurposing a selective inhibitor (epalrestat) that is already used clinically for diabetics.

## DISCUSSION

Accurate prediction of personalized drug responses is essential to reduce the persistently high recurrence rates following surgical resection of non-metastatic NSCLC (1, 2, 3). These high rates of locoregional and metastatic recurrences are due to drug resistant micro- and macrometastatic disease leading to treatment failure (46). Lack of high-throughput drug screening platforms is a clinical gap to improve NSCLC patient outcomes. Recent advancements in organoid technologies offer efficient platforms for personalized drug screening in cancer patients (13, 20, 47). In the present study, we generated PDTOs from treatment-naïve NSCLC patients undergoing curative surgical resection for non-metastatic disease. We observed PDTOs to be robust, quick, viable tumor-derived models that conserve histopathological features, genomic fidelity, and cellular signatures that are close to the matched patient primary lung tumor tissues. PDTO growth was accomplished from all ten NSCLC patients enrolled in our study. This high success rate is attributed to immediate processing (within 15 min of resection) of tumor specimens which maintains higher number of viable tumor cells. In addition, a lung cancer PDTO generation methodology was applied that was previously described by Li and colleagues with a success rate of >80% (26). All our PDTOs conserved histomorphology and diagnostic pathological biomarker expressions that were subtype-specific for LUAD and LUSC, respectively. Beyond these pathological evaluations applied in routine clinical care, recent studies revealed that viable PDTOs maintain additional features of the cancers from which they were derived over multiple time points, including tumor heterogeneity, genetic alterations, metabolism, and drug and radiation response (21, 22, 23, 24, 30). A critical component in the pre-clinical validation of tumor-derived models is the need for sustained conservation of complex genetic polymorphisms and mutational heterogeneity, cellular, molecular, and other biological properties of the matched parental tumors (48). In our cohort we demonstrate that the cellular, mutational landscape and molecular/transcriptomic phenotypes in NSCLC PDTOs are highly representative of the matched patient tumors, similar to reports by other groups (47). Whole exome sequencing confirmed that somatic mutations vastly matched between primary tumors and PDTOs, and only one out of six PDTOs showed a difference in a driver mutation (*KRAS*) in comparison to the primary tumor. Also, identified somatic mutations matched the ones commonly found in rural Midwest American cohorts consisting of heavy smokers (which is the same population we enrolled from for the present study) (49). Organoid platforms have also been utilized to understand the changes in transcriptional programs at various stages of tumorigenesis (50). Our findings confirm that PDTOs recapitulate the transcriptomic signature of matched patient tumors, also specifically with respect to epithelial (cancer) cell heterogeneity. LUAD-and LUSC-derived PDTOs showed abundance of AT2 cells and basal cells, respectively, that are both cancer histology-specific epithelial subtypes contributing to the onset and progression of LUAD/LUSC. Better understanding of the role of various cell types in NSCLC progression is needed, and investigators have used alveolar epithelial organoids as platforms to recapitulate early-stage LUAC progression (50).

Cancer PDTOs serve as suitable *ex vivo* platforms to evaluate drug responses in matched patients similar and possibly superior to testing *in vivo* PDX models (24, 51, 52). Here, we have performed drug testing in PDTOs derived from resected NSCLC tumors, frozen tumor tissues, and frozen PDTOs (living biobanks). We accomplished 100% PDTO reconstitution from a cryopreserved living biobank that confirms reports by other groups that achieved high success rates (>70%) of revival of PDTOs (51). While the utility of a living PDTO biobank will need to be proven by matching drug responses observed in patients to the ones observed in revived PDTOs, these living biobanks may provide highly valuable methods at later time points to determine future drug responses, if clinically needed. We applied a dynamic, real-time 3D growth analysis method via bright-field z-stack imaging of PDTOs to measure the real-time growth rates and the impact of drugs on PDTO growth over 6 days (28, 29). Although there are various adjuvant platinum-based doublet chemotherapy regimens administered to NSCLC patients that can be considered per treatment guidelines, for consistency and comparability we treated all PDTOs with a commonly given regimen of carboplatin/paclitaxel (53). We observed a drug response rate of 50% to chemotherapy treatments in the ten NSCLC PDTOs tested - similar to the ∼50% response rate observed in more advanced NSCLC patients undergoing chemotherapy treatments with platinum-based agents with radiographically detectable disease (3, 6). As the radiographically visible NSCLC disease was resected in our surgical patients at the time of administration of adjuvant chemotherapy, we were not able to directly compare response rates observed in PDTOs to the matched patients. However, one patient developed locoregional mediastinal lymph node recurrences and brain metastases under adjuvant chemotherapy with carboplatin/paclitaxel. Consistent with the assumed drug resistance to carboplatin/paclitaxel observed in this patient, the PDTO derived from this patient was also resistant towards carboplatin/paclitaxel. This observation is in alignment with other reports that demonstrated that the drug responses in PDTOs match the drug responses observed in NSCLC patients (19). These findings substantiate the clinical utility of PDTOs as reliable and high-throughput drug testing platforms in a clinically applicable timeframe.

Blockade of pro-tumorigenic mechanisms and pathways can lead to increased drug sensitivity, resulting in potentiation of therapeutic efficacy of chemotherapeutic drugs [41, 50]. To identify druggable targets, we screened PDTOs by whole transcriptome differential gene expression analysis. Aldo-keto reductase AKR1B10 was upregulated in chemoresistant NSCLC PDTOs. AKR1B10 overexpression has been reported in some solid tumors, including NSCLC (54). Moreover, the AKR1B family of enzymes are known for their role in detoxifying platinum-and taxol-based compounds causing them to be ineffective (55, 56). The AKR1B10 inhibitor epalrestat is a drug approved in Japan for patients with diabetic neuropathy (57). In America, there is an ongoing clinical trial to evaluate epalrestat in pediatric patients with Phosphomannomutase 2-congenital Disorder of Glycosylation (PMM2-CDG) (*ClinicalTrials.gov* Identifier: NCT04925960) (58). Following confirmation of AKR1B10 overexpression at the transcriptomic and protein level in chemoresistant PDTOs, we then repurposed epalrestat and observed conversion of chemoresistant PDTOs to chemosensitive with the addition of epalrestat. Importantly, epalrestat can be given orally and has penetration into the brain (59) as a common site of NSCLC metastases. Epalrestat has already been shown to overcome EGFR-targeted drug resistance in lung cancer cell lines and PDX mice (60). But beyond demonstrating the effect of epalrestat in overcoming chemoresistance, our study adds to the proof-of-concept that PDTOs are cost-and time-efficient testing platforms for repurposed drugs (12, 61). Of note, our group has developed explainable artificial intelligence algorithms to identify drug repurposing candidates in cancer patients by computational analysis of large-scale clinical and multiomics data (62). In future, repurposed drug candidates identified by machine learning or other means could be efficiently validated for cancer patients using PDTOs.

One of the limitations of our study is the small sample size of ten NSCLC patients only. While comprehensive PDTO drug testing and molecular characterization including the primary lung tumor tissue matching was performed, in future findings will need to be validated in a larger patient cohort. The resected surgical specimens that we processed delivered high tumor tissue quantities, likely enabling higher success rates for PDTO growth. Investigators have already developed PDTOs from low quantities of tumor tissue derived from percutaneous needle biopsies (63, 64). In surgically resected NSCLC patients, adjuvant systemic treatments targeting micrometastatic disease are associated with another dilemma: the inability to directly determine the effectiveness of a treatment response due to absence of radiographically detectable disease (65). We exclusively enrolled NSCLC patients undergoing immediate surgical resection of their tumor disease. At least, we could successfully match the chemoresponse observed in a PDTO in one patient who developed progressive disease under carboplatin/paclitaxel. The remaining nine study patients remain recurrence-free during the observation period of more than two years and are being monitored closely during their follow-up visits.

Then, limited or lack of preservation of the immune system and other microenvironmental cells may have critical impact on treatment responses observed in PDTOs and limits PDTO testing involving immunotherapeutic strategies. Although 3D culture systems mimic the microenvironment, PDTOs still often grow as rather pure populations of cancer cells while other microenvironmental cells get lost in the culturing process (66). To overcome this general limitation of PDTOs, in future co-culture systems involving PDTOs and matched tumor-infiltrating lymphocytes or peripheral blood mononuclear cells will allow to determine the impact of the local tumor immune microenvironment or the systemic immune status in accurate prediction of patient responses to treatments, including immunotherapies (67).

## CONCLUSIONS

Personalized drug response testing can be performed within a clinically applicable time in non-metastatic, surgically treated NSCLC patients by generating PDTOs, including from a frozen living biobank. The use of PDTOs to identify mutational landscape and tumor heterogeneity of the matched patient tumor to potentially predict the clonal evolution of the patient’s cancer in response to therapies provides highly valuable information to design personal precision therapies for NSCLC patients. The time-and cost-efficient approach with viable PDTOs is an exciting methodology to improve NSCLC patient outcomes by avoiding unnecessary and toxic therapies while allowing for the escalation of effective therapies in patients to prevent and/or treat recurrences and reduce overall mortality of this most devastating cancer.

### LIST OF ABBREVIATIONS

AT2 cells: alveolar epithelial cell type II
AJCC: American Joint Committee on Cancer
ANOVA: analysis of variance
adDMEM: advanced Dulbecco’s Modified Eagle Medium
CK: cytokeratin
DAB: diaminobenzidine
DMSO: dimethyl sulfoxide
DNA: deoxyribonucleic acid
EGF: epidermal growth factor
EGFR: epidermal growth factor receptor
FBS: fetal bovine serum
FGF: fibroblast growth factor
H&E: hematoxylin & eosin
HRP: horseradish peroxidase
IASLC: International Association for the Study of Lung Cancer
IHC: immunohistochemistry
IRB: Institutional Review Board
LGEA: lung gene expression analysis
LUAC: lung adenocarcinoma
LUSC: lung squamous cell carcinoma
N/A: not applicable
NSCLC: non-small cell lung cancer
PCA: principal component analysis
PCR: polymerase chain reaction
PDTO: patient-derived tumor organoid
PDX: patient-derived xenograft
RNA: ribonucleic acid
TNM: tumor node metastasis
TTF-1: thyroid transcription factor
UICC: Union for International Cancer Control (UICC)
WES: whole exome sequencing

## DECLARATIONS

### Ethics approval and consent to participate

The Institutional Review Board (IRB) of the University of Missouri (MU) approved this study (IRB#: 2010166). Trials were registered at *ClinicalTrials.gov* (Trial registration number (TRN): identifier NCT02838836; date of registration: July 20, 2016).

### Consent for publication

All patients gave written informed consent. Availability of data and materials: All data generated or analyzed during this study, if not included in this article and its supplementary information files, are available from the corresponding authors on reasonable request.

### Availability of data and material

All raw whole exome and RNA sequencing data were deposited in NCBI Sequence Read Archive (SRA) and are accessible through project numbers PRJNA1020294 and PRJNA935652, respectively. All other datasets generated and analyzed in the current study are available from the corresponding author upon request.

### Competing interests

The authors declare no competing interests.

## Funding

This study was supported by an Ellis Fischel Cancer Center Pilot Award (J.T.K.). J.B.M. received funding from the Department of Veterans Affairs K2BX004346-01A1. M.A.C has support from the Washington University DDRCC (NIDDK P30 DK052574) and Barnes-Jewish Hospital Foundation Siteman Investment Program Award, Grant 5897. The content is solely the responsibility of the authors and does not necessarily represent the official views of the Department of Veterans Affairs. The funding bodies had no role in study design, collection, analysis, interpretation of data or writing the manuscript. This research project was also supported by Endowments from the University of Missouri [Paul K. and Dianne Shumaker (C.R.S.) and Margaret Proctor Mulligan (J.T.K.) Professor Endowments].

### Author contributions

K.N.S., Y.M., M.G., J.T.K.: enrolled the patients and conducted the experiments. Y.I.N., K.N.S., J.B.M., W.C.W., S.R., C.R.S.: analyzed whole exome and transcriptomics data, W.C.W., J.B.M., M.A.C., S.R., J.T.K.: supervised data analysis. K.N.S., Y.M., A.S., G.L., M.A.C., J.B.M., S.R., J.T.K.: conceptualization, methodology, formal analysis, original draft, writing – review & editing. Y.M., J.T.K., F.G.: Statistical analyses.

All authors reviewed the manuscript.

## Acknowledgements

We are exceedingly grateful to all patients for their voluntary participation. The authors thank Nathan Bivens (MU Genomics Technology Core) for technical support.

## SUPPLEMENTARY TABLE

**Supplementary Table 1:** NSCLC Tier 1 SNPs identified in patient primary tumors and matched PDTOs.

| Mutational details | Samples |  |  |  |  |  |
| --- | --- | --- | --- | --- | --- | --- |
|  | MU374 | MU377 | MU378 | MU383 | MU387 | MU388 |
| **Gene: EGFR**<br>Chromosome: 7<br>Exon: exon13<br>CDS Mutation: c.G1562A<br>AA mutation: R521K<br>Cosmic ID: COSM3721608 | PT:<br>Found<br>PDTO:<br>Found | PT:<br>Found<br>PDTO:<br>Found | PT:<br>Not Found<br>PDTO: Not<br>Found | PT:<br>Not Found<br>PDTO:<br>Not Found | PT:<br>Not Found<br>PDTO: Not<br>Found | PT:<br>Found<br>PDTO:<br>Found |
| **Gene: TP53**<br>Chromosome: 17<br>Exon: exon4<br>CDS Mutation: c.C215G<br>AA mutation: P72R<br>Cosmic ID: COSM250061 | PT:<br>Found<br>PDTO:<br>Found | PT:<br>Not Found<br>PDTO:<br>Not Found | PT:<br>Found<br>PDTO:<br>Found | PT:<br>Found<br>PDTO:<br>Found | PT:<br>Found<br>PDTO:<br>Found | PT:<br>Found<br>PDTO:<br>Found |
| **Gene: TP53**<br>Chromosome: 17<br>Exon: exon5<br>CDS Mutation: c.G517A<br>AA mutation: V173M<br>Cosmic ID: COSM98966 | PT:<br>Not Found<br>PDTO:<br>Not Found | PT:<br>Not Found<br>PDTO:<br>Not Found | PT:<br>Not Found<br>PDTO:<br>Not Found | PT:<br>Not Found<br>PDTO:<br>Not Found | PT:<br>Not Found<br>PDTO:<br>Not Found | PT:<br>Found<br>PDTO:<br>Not Found |
| **Gene: KRAS**<br>Chromosome: 12<br>Exon: exon2<br>CDS Mutation: c.G35T<br>AA mutation: G12V<br>Cosmic ID: COSM1140133 | PT:<br>Found<br>PDTO:<br>Not Found | PT:<br>Not Found<br>PDTO: Not<br>Found | PT:<br>Not Found<br>PDTO:<br>Not Found | PT:<br>Not Found<br>PDTO:<br>Not Found | PT:<br>Not Found<br>PDTO:<br>Not Found | PT:<br>Not Found<br>PDTO:<br>Not Found |
| **Gene: ALK**<br>Chromosome: 2<br>Exon: exon29<br>CDS Mutation: c.C4587G<br>AA mutation: D1529E<br>Cosmic ID: COSM3758201 | PT:<br>Not Found<br>PDTO:<br>Not Found | PT:<br>Not Found<br>PDTO:<br>Not Found | PT:<br>Not Found<br>PDTO:<br>Not Found | PT:<br>Not Found<br>PDTO:<br>Not Found | PT:<br>Not Found<br>PDTO:<br>Not Found | PT:<br>Found<br>PDTO:<br>Found |
| **Gene: KDR**<br>Chromosome: 4<br>Exon: exon11<br>CDS Mutation: c.A1416T<br>AA mutation: Q472H<br>Cosmic ID: COSM149673 | PT:<br>Not Found<br>PDTO:<br>Not Found | PT:<br>Found<br>PDTO:<br>Found | PT:<br>Not Found<br>PDTO:<br>Not Found | PT:<br>Not Found<br>PDTO:<br>Not Found | PT:<br>Not Found<br>PDTO:<br>Not Found | PT:<br>Found<br>PDTO:<br>Found |
| **Gene: KDR**<br>Chromosome: 4<br>Exon: exon7<br>CDS Mutation: c.G889A<br>AA mutation: V297I<br>Cosmic ID: COSM1131107 | PT:<br>Not Found<br>PDTO:<br>Not Found | PT:<br>Not Found<br>PDTO:<br>Not Found | PT:<br>Not Found<br>PDTO:<br>Not Found | PT:<br>Not Found<br>PDTO:<br>Not Found | PT:<br>Not Found<br>PDTO:<br>Not Found | PT:<br>Found<br>PDTO:<br>Found |
| **Gene: RET**<br>Chromosome: 10<br>Exon: exon11<br>CDS Mutation: c.G2071A<br>AA mutation: G691S<br>Cosmic ID: COSM1666596 | PT:<br>Not Found<br>PDTO:<br>Not Found | PT:<br>Not Found<br>PDTO:<br>Not Found | PT:<br>Not Found<br>PDTO:<br>Not Found | PT:<br>Not Found<br>PDTO:<br>Not Found | PT:<br>Found<br>PDTO:<br>Found | PT:<br>Not Found<br>PDTO:<br>Not Found |

PT: Primary Tumor, PDTO: Patient-derived tumor organoid.

## SUPPLEMENTARY FIGURES AND LEGENDS

**Supplementary Figure 1.**
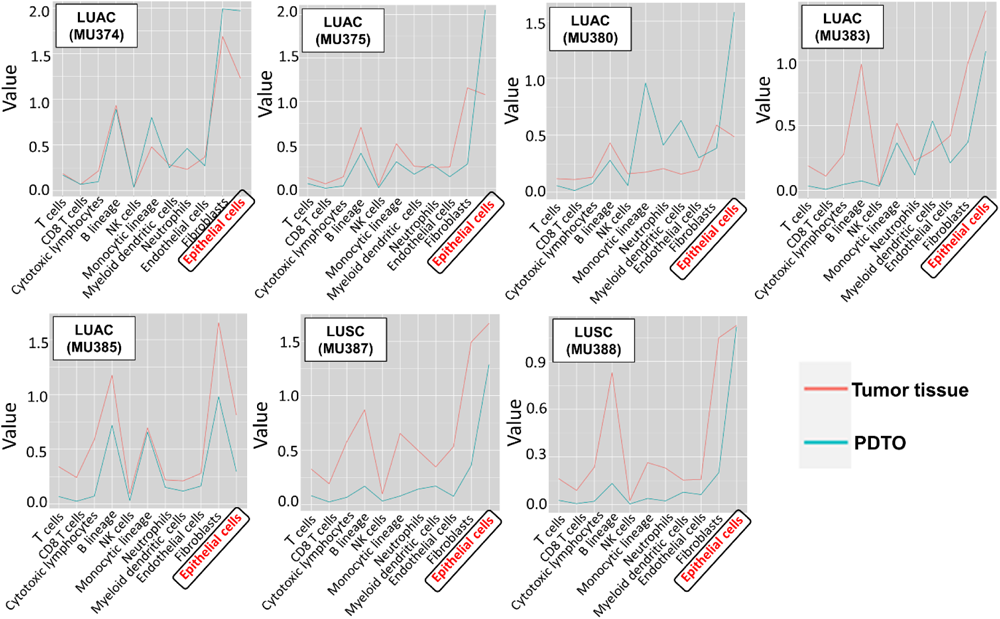
Cellular heterogeneity comparison between matched PDTOs and patient primary tumors (PT). Microenvironment Cell Populations (MCP)-counter graphs showing cellular abundance of matched PTs and PDTOs of all NSCLC patients with conserved epithelial cell populations, whereas immune, stromal, and endothelial cell compositions were found to be not conserved in all samples except MU385.

**Supplementary Figure 2.**
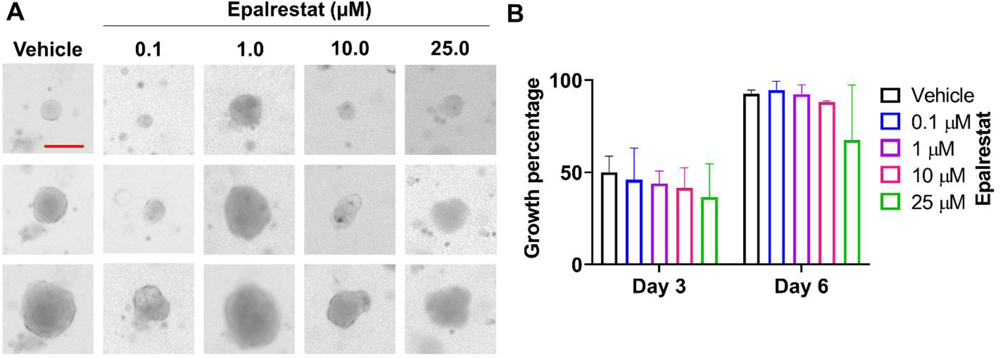
Dose titration of epalrestat against PDTOs to determine the cytotoxicity. To determine the dose for drug repurposing experiments, epalrestat was titrated against PDTOs in increasing micro molar concentrations and growth measurements was done against vehicle control having day 0 as baseline. **A:** Representative image of PDTOs from vehicle and treatment groups, scale bar 200 µm. **B:** Graph showing accumulated results of dose titration experiment where epalrestat was not found to be cytotoxic up to 25 µM concentration.

## Notes

### Competing Interest Statement

The authors have declared no competing interest.

## REFERENCES

1. Siegel RL, Miller KD, Fuchs HE, Jemal A. Cancer statistics, 2022. CA Cancer J Clin. 2022;72(1):7-33.

2. Molina JR, Yang P, Cassivi SD, Schild SE, Adjei AA. Non-small cell lung cancer: epidemiology, risk factors, treatment, and survivorship. Mayo Clin Proc. 2008;83(5):584–94.

3. Arriagada R, Bergman B, Dunant A, Le Chevalier T, Pignon JP, Vansteenkiste J, et al. Cisplatin-based adjuvant chemotherapy in patients with completely resected non-small-cell lung cancer. N Engl J Med. 2004;350(4):351–60.

4. Krebs MG, Sloane R, Priest L, Lancashire L, Hou JM, Greystoke A, et al. Evaluation and prognostic significance of circulating tumor cells in patients with non-small-cell lung cancer. J Clin Oncol. 2011;29(12):1556–63.

5. Pantel K, Izbicki J, Passlick B, Angstwurm M, Haussinger K, Thetter O, et al. Frequency and prognostic significance of isolated tumour cells in bone marrow of patients with non-small-cell lung cancer without overt metastases. Lancet. 1996;347(9002):649-53.

6. Pisters KM, Vallieres E, Crowley JJ, Franklin WA, Bunn PA, Jr., Ginsberg RJ, et al. Surgery with or without preoperative paclitaxel and carboplatin in early-stage non-small-cell lung cancer: Southwest Oncology Group Trial S9900, an intergroup, randomized, phase III trial. J Clin Oncol. 2010;28(11):1843–9.

7. Wu YL, Tsuboi M, He J, John T, Grohe C, Majem M, et al. Osimertinib in Resected EGFR-Mutated Non-Small-Cell Lung Cancer. N Engl J Med. 2020;383(18):1711–23.

8. O’Brien M, Paz-Ares L, Marreaud S, Dafni U, Oselin K, Havel L, et al. Pembrolizumab versus placebo as adjuvant therapy for completely resected stage IB-IIIA non-small-cell lung cancer (PEARLS/KEYNOTE-091): an interim analysis of a randomised, triple-blind, phase 3 trial. Lancet Oncol. 2022;23(10):1274–86.

9. Forde PM, Spicer J, Lu S, Provencio M, Mitsudomi T, Awad MM, et al. Neoadjuvant Nivolumab plus Chemotherapy in Resectable Lung Cancer. N Engl J Med. 2022;386(21):1973–85.

10. Fountzilas E, Tsimberidou AM. Overview of precision oncology trials: challenges and opportunities. Expert Rev Clin Pharmacol. 2018;11(8):797–804.

11. Mok TSK, Wu YL, Kudaba I, Kowalski DM, Cho BC, Turna HZ, et al. Pembrolizumab versus chemotherapy for previously untreated, PD-L1-expressing, locally advanced or metastatic non-small-cell lung cancer (KEYNOTE-042): a randomised, open-label, controlled, phase 3 trial. Lancet. 2019;393(10183):1819-30.

12. Jin G, Wong ST. Toward better drug repositioning: prioritizing and integrating existing methods into efficient pipelines. Drug Discov Today. 2014;19(5):637–44.

13. Jenkins RW, Aref AR, Lizotte PH, Ivanova E, Stinson S, Zhou CW, et al. Ex Vivo Profiling of PD-1 Blockade Using Organotypic Tumor Spheroids. Cancer Discov. 2018;8(2):196–215.

14. Lallo A, Schenk MW, Frese KK, Blackhall F, Dive C. Circulating tumor cells and CDX models as a tool for preclinical drug development. Transl Lung Cancer Res. 2017;6(4):397–408.

15. Suvilesh KN, Nussbaum YI, Radhakrishnan V, Manjunath Y, Avella DM, Staveley-O’Carroll KF, et al. Tumorigenic circulating tumor cells from xenograft mouse models of non-metastatic NSCLC patients reveal distinct single cell heterogeneity and drug responses. Mol Cancer. 2022;21(1):73.

16. Suvilesh KN, Manjunath Y, Pantel K, Kaifi JT. Preclinical models to study patient-derived circulating tumor cells and metastasis. Trends Cancer. 2023;9(4):355–71.

17. Tentler JJ, Tan AC, Weekes CD, Jimeno A, Leong S, Pitts TM, et al. Patient-derived tumour xenografts as models for oncology drug development. Nat Rev Clin Oncol. 2012;9(6):338–50.

18. Suvilesh KN, Manjunath Y, Pantel K, Kaifi JT. Preclinical models to study patient-derived circulating tumor cells and metastasis. Trends Cancer. 2023.

19. Shi R, Radulovich N, Ng C, Liu N, Notsuda H, Cabanero M, et al. Organoid Cultures as Preclinical Models of Non-Small Cell Lung Cancer. Clin Cancer Res. 2020;26(5):1162–74.

20. Tuveson D, Clevers H. Cancer modeling meets human organoid technology. Science. 2019;364(6444):952-5.

21. Boj SF, Hwang CI, Baker LA, Chio, II, Engle DD, Corbo V, et al. Organoid models of human and mouse ductal pancreatic cancer. Cell. 2015;160(1-2):324–38.

22. Vlachogiannis G, Hedayat S, Vatsiou A, Jamin Y, Fernandez-Mateos J, Khan K, et al. Patient-derived organoids model treatment response of metastatic gastrointestinal cancers. Science. 2018;359(6378):920-6.

23. van de Wetering M, Francies HE, Francis JM, Bounova G, Iorio F, Pronk A, et al. Prospective derivation of a living organoid biobank of colorectal cancer patients. Cell. 2015;161(4):933–45.

24. Pasch CA, Favreau PF, Yueh AE, Babiarz CP, Gillette AA, Sharick JT, et al. Patient-Derived Cancer Organoid Cultures to Predict Sensitivity to Chemotherapy and Radiation. Clin Cancer Res. 2019;25(17):5376–87.

25. Amin MB, Greene FL, Edge SB, Compton CC, Gershenwald JE, Brookland RK, et al. The Eighth Edition AJCC Cancer Staging Manual: Continuing to build a bridge from a population-based to a more “personalized” approach to cancer staging. CA Cancer J Clin. 2017;67(2):93–9.

26. Li Z, Yu L, Chen D, Meng Z, Chen W, Huang W. Protocol for generation of lung adenocarcinoma organoids from clinical samples. STAR Protoc. 2021;2(1):100239.

27. Manjunath Y, Upparahalli SV, Avella DM, Deroche CB, Kimchi ET, Staveley-O’Carroll KF, et al. PD-L1 Expression with Epithelial Mesenchymal Transition of Circulating Tumor Cells Is Associated with Poor Survival in Curatively Resected Non-Small Cell Lung Cancer. Cancers (Basel). 2019;11(6).

28. Walsh AJ, Poole KM, Duvall CL, Skala MC. Ex vivo optical metabolic measurements from cultured tissue reflect in vivo tissue status. J Biomed Opt. 2012;17(11):116015.

29. Walsh AJ, Skala MC. Optical metabolic imaging quantifies heterogeneous cell populations. Biomed Opt Express. 2015;6(2):559–73.

30. Chen W, Wong C, Vosburgh E, Levine AJ, Foran DJ, Xu EY. High-throughput image analysis of tumor spheroids: a user-friendly software application to measure the size of spheroids automatically and accurately. J Vis Exp. 2014(89).

31. Li H, Durbin R. Fast and accurate long-read alignment with Burrows-Wheeler transform. Bioinformatics. 2010;26(5):589–95.

32. Tarasov A, Vilella AJ, Cuppen E, Nijman IJ, Prins P. Sambamba: fast processing of NGS alignment formats. Bioinformatics. 2015;31(12):2032–4.

33. DePristo MA, Banks E, Poplin R, Garimella KV, Maguire JR, Hartl C, et al. A framework for variation discovery and genotyping using next-generation DNA sequencing data. Nat Genet. 2011;43(5):491–8.

34. Wang K, Li M, Hakonarson H. ANNOVAR: functional annotation of genetic variants from high-throughput sequencing data. Nucleic Acids Res. 2010;38(16):e164.

35. Tate JG, Bamford S, Jubb HC, Sondka Z, Beare DM, Bindal N, et al. COSMIC: the Catalogue Of Somatic Mutations In Cancer. Nucleic Acids Res. 2019;47(D1):D941–D7.

36. Sondka Z, Bamford S, Cole CG, Ward SA, Dunham I, Forbes SA. The COSMIC Cancer Gene Census: describing genetic dysfunction across all human cancers. Nat Rev Cancer. 2018;18(11):696–705.

37. Ewels P, Magnusson M, Lundin S, Kaller M. MultiQC: summarize analysis results for multiple tools and samples in a single report. Bioinformatics. 2016;32(19):3047–8.

38. Dobin A, Davis CA, Schlesinger F, Drenkow J, Zaleski C, Jha S, et al. STAR: ultrafast universal RNA-seq aligner. Bioinformatics. 2013;29(1):15–21.

39. Liao Y, Smyth GK, Shi W. featureCounts: an efficient general purpose program for assigning sequence reads to genomic features. Bioinformatics. 2014;30(7):923–30.

40. Love MI, Huber W, Anders S. Moderated estimation of fold change and dispersion for RNA-seq data with DESeq2. Genome Biol. 2014;15(12):550.

41. Becht E, Giraldo NA, Lacroix L, Buttard B, Elarouci N, Petitprez F, et al. Estimating the population abundance of tissue-infiltrating immune and stromal cell populations using gene expression. Genome Biol. 2016;17(1):218.

42. Du Y, Kitzmiller JA, Sridharan A, Perl AK, Bridges JP, Misra RS, et al. Lung Gene Expression Analysis (LGEA): an integrative web portal for comprehensive gene expression data analysis in lung development. Thorax. 2017;72(5):481–4.

43. Karlsson M, Zhang C, Mear L, Zhong W, Digre A, Katona B, et al. A single-cell type transcriptomics map of human tissues. Sci Adv. 2021;7(31).

44. Zhang X, Lan Y, Xu J, Quan F, Zhao E, Deng C, et al. CellMarker: a manually curated resource of cell markers in human and mouse. Nucleic Acids Res. 2019;47(D1):D721–D8.

45. Wilkerson MD, Schallheim JM, Hayes DN, Roberts PJ, Bastien RR, Mullins M, et al. Prediction of lung cancer histological types by RT-qPCR gene expression in FFPE specimens. J Mol Diagn. 2013;15(4):485–97.

46. Sosa Iglesias V, Giuranno L, Dubois LJ, Theys J, Vooijs M. Drug Resistance in Non-Small Cell Lung Cancer: A Potential for NOTCH Targeting? Front Oncol. 2018;8:267.

47. Li Z, Qian Y, Li W, Liu L, Yu L, Liu X, et al. Human Lung Adenocarcinoma-Derived Organoid Models for Drug Screening. iScience. 2020;23(8):101411.

48. Schutte M, Risch T, Abdavi-Azar N, Boehnke K, Schumacher D, Keil M, et al. Molecular dissection of colorectal cancer in pre-clinical models identifies biomarkers predicting sensitivity to EGFR inhibitors. Nat Commun. 2017;8:14262.

49. Mitchem JB, Miller A, Manjunath Y, Barbirou M, Raju M, Shen Y, et al. Somatic mutation variant analysis in rural, resectable non-small cell lung carcinoma patients. Cancer Genet. 2022;268–269:75-82.

50. Dost AFM, Moye AL, Vedaie M, Tran LM, Fung E, Heinze D, et al. Organoids Model Transcriptional Hallmarks of Oncogenic KRAS Activation in Lung Epithelial Progenitor Cells. Cell Stem Cell. 2020;27(4):663–78 e8.

51. Kim M, Mun H, Sung CO, Cho EJ, Jeon HJ, Chun SM, et al. Patient-derived lung cancer organoids as in vitro cancer models for therapeutic screening. Nat Commun. 2019;10(1):3991.

52. Hu Y, Sui X, Song F, Li Y, Li K, Chen Z, et al. Lung cancer organoids analyzed on microwell arrays predict drug responses of patients within a week. Nat Commun. 2021;12(1):2581.

53. Strauss GM, Herndon JE, 2nd, Maddaus MA, Johnstone DW, Johnson EA, Harpole DH, et al. Adjuvant paclitaxel plus carboplatin compared with observation in stage IB non-small-cell lung cancer: CALGB 9633 with the Cancer and Leukemia Group B, Radiation Therapy Oncology Group, and North Central Cancer Treatment Group Study Groups. J Clin Oncol. 2008;26(31):5043-51.

54. Fukumoto S, Yamauchi N, Moriguchi H, Hippo Y, Watanabe A, Shibahara J, et al. Overexpression of the aldo-keto reductase family protein AKR1B10 is highly correlated with smokers’ non-small cell lung carcinomas. Clin Cancer Res. 2005;11(5):1776–85.

55. Matsunaga T, Wada Y, Endo S, Soda M, El-Kabbani O, Hara A. Aldo-Keto Reductase 1B10 and Its Role in Proliferation Capacity of Drug-Resistant Cancers. Front Pharmacol. 2012;3:5.

56. Penning TM, Jonnalagadda S, Trippier PC, Rizner TL. Aldo-Keto Reductases and Cancer Drug Resistance. Pharmacol Rev. 2021;73(3):1150–71.

57. Hotta N, Akanuma Y, Kawamori R, Matsuoka K, Oka Y, Shichiri M, et al. Long-term clinical effects of epalrestat, an aldose reductase inhibitor, on diabetic peripheral neuropathy: the 3-year, multicenter, comparative Aldose Reductase Inhibitor-Diabetes Complications Trial. Diabetes Care. 2006;29(7):1538–44.

58. Iyer S, Sam FS, DiPrimio N, Preston G, Verheijen J, Murthy K, et al. Repurposing the aldose reductase inhibitor and diabetic neuropathy drug epalrestat for the congenital disorder of glycosylation PMM2-CDG. Dis Model Mech. 2019;12(11).

59. Ramirez MA, Borja NL. Epalrestat: an aldose reductase inhibitor for the treatment of diabetic neuropathy. Pharmacotherapy. 2008;28(5):646–55.

60. Zhang KR, Zhang YF, Lei HM, Tang YB, Ma CS, Lv QM, et al. Targeting AKR1B1 inhibits glutathione de novo synthesis to overcome acquired resistance to EGFR-targeted therapy in lung cancer. Sci Transl Med. 2021;13(614):eabg6428.

61. Hirt CK, Booij TH, Grob L, Simmler P, Toussaint NC, Keller D, et al. Drug screening and genome editing in human pancreatic cancer organoids identifies drug-gene interactions and candidates for off-label treatment. Cell Genom. 2022;2(2):100095.

62. Al-Taie Z, Liu D, Mitchem JB, Papageorgiou C, Kaifi JT, Warren WC, et al. Explainable artificial intelligence in high-throughput drug repositioning for subgroup stratifications with interventionable potential. J Biomed Inform. 2021;118:103792.

63. Choi SI, Jeon AR, Kim MK, Lee YS, Im JE, Koh JW, et al. Development of Patient-Derived Preclinical Platform for Metastatic Pancreatic Cancer: PDOX and a Subsequent Organoid Model System Using Percutaneous Biopsy Samples. Front Oncol. 2019;9:875.

64. Vilgelm AE, Bergdorf K, Wolf M, Bharti V, Shattuck-Brandt R, Blevins A, et al. Fine-Needle Aspiration-Based Patient-Derived Cancer Organoids. iScience. 2020;23(8):101408.

65. Dasari A, Grothey A, Kopetz S. Circulating Tumor DNA-Defined Minimal Residual Disease in Solid Tumors: Opportunities to Accelerate the Development of Adjuvant Therapies. J Clin Oncol. 2018;36(35):JCO2018789032.

66. Dijkstra KK, Monkhorst K, Schipper LJ, Hartemink KJ, Smit EF, Kaing S, et al. Challenges in Establishing Pure Lung Cancer Organoids Limit Their Utility for Personalized Medicine. Cell Rep. 2020;31(5):107588.

67. Cattaneo CM, Dijkstra KK, Fanchi LF, Kelderman S, Kaing S, van Rooij N, et al. Tumor organoid-T-cell coculture systems. Nat Protoc. 2020;15(1):15–39.

